# Early detection of oxidative stress in mammary epithelial cells by using secreted Cyclophilin A as a marker in the milk

**DOI:** 10.1101/2025.07.29.667364

**Authors:** Sravani Kavati, Geethu Krishna, Ilsha Pathan, Khushi Jatav, Goutam Ulgekar, Nirmalya Ganguli, Subeer S Majumdar, Surya Ramachandran

## Abstract

**Objective:** To investigate the secretion and expression of cyclophilin A (CypA) in mammary epithelial cells *in vitro* under oxidative stress conditions and to evaluate its potential as a biomarker of oxidative stress in dairy cows.

**Methods:** Primary mammary epithelial cells isolated from goat mammary gland, luminal epithelial cells isolated from goat milk and mouse mammary epithelial cells were cultured and treated with hydrogen peroxide (H_2_O_2_) and tert-butyl hydroperoxide (t-BHP) to induce oxidative stress. Cell viability, intracellular reactive oxygen species (ROS) levels, lipid peroxidation, and antioxidant enzyme activity were measured to assess the oxidative status of cells. CypA gene expression was quantified using quantitative reverse transcription polymerase chain reaction and the secreted CypA levels were measured by using enzyme linked immunosorbent assay and immunoblotting. Milk samples from various cow breeds were also analysed for somatic cell count, catalase activity and CypA levels.

**Results:** The primary mammary epithelial cells culture displayed typical epithelial cell morphology. Oxidative stress induced by 200 µM of both H_2_O_2_ and t-BHP and an exposure duration of 1 hour was observed to be optimum in inducing oxidative stress. A significant increase in ROS levels and lipid peroxidation index was observed across all cell types. Catalase activity was also elevated in the treated cells. An upregulation of CypA gene expression was observed in response to oxidative stress with a significant increase in CypA protein levels. Milk samples from the various cattle breeds revealed varying somatic cell counts with the highest in Kankrej and lowest in Holstein Friesian.

**Conclusion:** These findings demonstrate that CypA is secreted from the mammary epithelial cells in response to oxidative stress and may serve as a potential biomarker for oxidative stress in dairy cows.

## INTRODUCTION

Oxidative stress is a phenomenon in which there is a disequilibrium between pro-oxidants and antioxidants [1]. It involves increased production of free radical oxidants such as reactive oxygen species (ROS) that are capable of oxidizing macromolecules including membrane lipids, structural proteins, nucleic acids, and enzymes and enzymes which may interfere with the normal cell function and can be detrimental to cell [2]. Oxidative stress has been implicated in a wide range of pathologies as their primary cause and is known to promote disease progression [3]. A compromised oxidative status is well known to cause the immunocompetence in animals and affects overall livestock health, efficiency and primarily milk production [4].

ROS are known to trigger various cellular signalling pathways associated with cell survival, repair, cell death and inflammation [5]. Cyclophilins are a family of proteins that belong to immunophilins and are secreted from the cells especially vascular smooth muscle cells and endothelial cells, in response to ROS levels. They are characterised by peptidyl-prolyl cis-trans isomerase activity at proline residues [6]. Cyclophilin A (CypA) is the most abundant among cyclophilins and is reported to have important roles in many biological conditions, including protein folding, trafficking, and T-cell activation. Extracellular secretion of CypA from endothelial cells and vascular smooth muscle cells occurs in response to inflammatory stimuli, including hypoxia, infection, and oxidative stress [7]. A recent study has shown the chemotactic role of CypA in inducing inflammation under conditions of mastitis in dairy cows [8].

Dairy cows may encounter increased oxidative stress and disease susceptibility during the various stages of their productive period due to several reasons [9]. In dairy cows, easily accessible oxidative stress biomarkers may serve as a useful tool to detect the oxidative status thereby allowing to adopt possible strategies to restrain the negative effects of free radicals on animal’s health. Routine serum biochemical analyses are a reliable way to understand the health status of animal [10]. Commonly assessed stress-related biomarkers from blood are the levels of antioxidants such as superoxide dismutase, glutathione peroxidase, catalase activity etc. [11]. Given the prevalence of oxidative stress-related health complications, there is a need for a definitive biomarker to detect and quantify the level of actual oxidative stress in dairy cows in a non-invasive manner.

In this study, we have developed an *in vitro* cell model using mammary epithelial cells of goat and mouse origin by inducing oxidative stress in these cells using hydrogen peroxide (H_2_O_2_) and tert-butyl hydroperoxide (t-BHP). Here, we have demonstrated the secretion of CypA from these cells in response to induced oxidative stress.

## MATERIALS AND METHODS

### In vitro cellular model of mammary epithelial cells

Three different cell types were used in the study; primary mammary epithelial cells isolated from goat mammary gland (GMECs), luminal epithelial cells isolated from goat milk (GLECs) and mouse mammary epithelial cells (MMECs; HC11 cell line; sourced from NIAB, Telengana, India). GMECs and GLECs were cultured using 90% Dulbecco’s Modified Eagle Medium (DMEM)/F-12 supplemented with 10% fetal bovine serum (FBS), and 1% antibiotic-antimycotic cocktail; insulin, transferrin, and selenium (ITS) supplement (5 μg/mL) and epidermal growth factor (EGF; 10 ng/mL). MMECs were cultured in 89% RPMI (Roswell Park Memorial Institute) 1640 medium supplemented with 10% FBS, and 1%, antibiotic-antimycotic cocktail. The cells were seeded in T-25 tissue culture flasks and incubated at 37ºC with 5% carbon dioxide (CO_2_) until the cells reached 70-80% confluency.

Trypan blue dye (0.4%) was used to stain the cells to check for viability. Viable cells (n=10,000) were seeded onto each well of a 96-well plate to facilitate cell adhesion, followed by the addition of DMEM/RPMI media to the cells in culture. The plate was incubated in a CO_2_ incubator at 37°C with 95% humidity for 24 hours for further experiments.

### Induction of oxidative stress

Hydrogen peroxide (H_2_O_2_) and tert-butyl hydrogen peroxide (t-BHP) were used to induce oxidative stress in GMEC, GLEC and MMEC cultures. After the incubation period of 24 hours, the cells were treated with 100µL of H_2_O_2_ and t-BHP in varying concentrations such as 100 μM, 200 μM, 300 μM, 400 μM and 500 μM, for a duration of 1 to 4 hours. Post-treatment, the cells were subjected to viability assays.

### Cell viability assay

The effect of H_2_O_2_ /t-BHP treatment on the viability of cells was determined using 3-(4, 5-dimethylthiazol-2-yl)-2,5-diphenyl tetrazolium bromide (MTT, Hi-media) assay. Briefly, the cells in 96-well microplates with each treatment group were washed with fresh culture medium and then incubated in fresh medium containing MTT (0.5 mg/mL) for 4 hours at 37°C. After which, the medium was discarded and the cells were incubated in dimethyl sulfoxide (DMSO, Sigma-Aldrich) to dissolve formazan aggregates produced during the reaction. The absorbance of the converted dye was measured at a wavelength of 570 nm using a microplate reader.

### Measurement of ROS levels

An oxidation-sensitive fluorescent dye, 2′,7′-dichlorodihydrofluorescein diacetate; (DCF-DA) was used to quantify the ROS levels in H_2_O_2_ /t-BHP treated and control cells. Briefly, the cells were incubated with 10 µM of DCF-DA (Sigma) at 37º in a CO_2_ incubator for 30 minutes. Light emitted by the cells was examined under a fluorescence microscope with an excitation wavelength of 485 nm and an emission wavelength of 530 nm. Cells stained with DAPI were used as a positive control.

### Measurement of the levels of lipid peroxidation

Thiobarbituric Acid Reactive Substances (TBARS) assay was used to assess the level of lipid peroxidation in cells using TBARS analysis kit (Abcam). The cells were washed with sterile phosphate-buffered saline (PBS) to remove any traces of media and oxidizing agents. Following cell lysis, the suspension was centrifuged, and the supernatant was used for performing the assay based on the kit instructions. Varying concentrations of malondialdehyde provided in the kit served as standards. The OD was read at a wavelength of 532 nm using a microplate reader.

### Catalase assay

The enzymatic activity of catalase to oxidize H_2_O_2_ was analysed using catalase assay. Cell lysates of the control and H_2_O_2_ /t-BHP treated cells were reacted with catalase enzyme (Sigma-Aldrich). Absorbance of the coloured product formed was measured at 405 nm. Catalase enzyme standards were used to plot the standard curve.

### Gene expression analysis of *cypA* by quantitative reverse transcription PCR (qRT-PCR)

Total cellular RNA was isolated from the oxidative stress-induced and control cells using an RNA extraction kit (Qiagen) following the manufacturer’s protocol. The RNA yield and purity were determined using a Nanodrop spectrophotometer. The total RNA was reverse transcribed into cDNA using a cDNA synthesis kit (Himedia) following the manufacturer’s instructions. The primers for *cypA* gene were designed using the Primer-BLAST software and are listed in Table 1. The qRT-PCR technique, along with the SYBR Green (Thermofisher) protocol, was utilized to quantify the expression of *cypA* gene. The experiment was performed using QuantStudio 7 Flex qRT-PCR System.

**Table 1:**
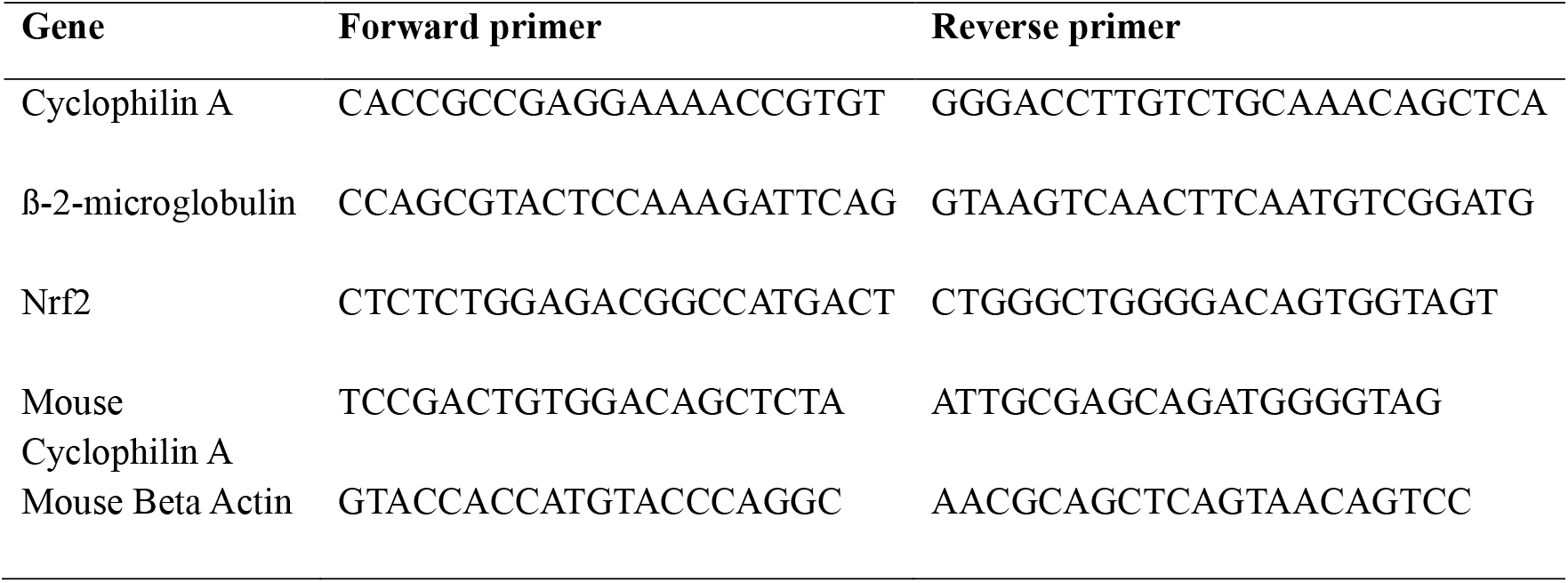
The sequences of primers used for RT-PCR.

### Measurement of CypA levels in the conditioned media

The levels of secreted CypA from supernatants of GMECs, GLECs, and MMECs cultures were quantified using ELISA. The cells after attaining 80% confluency were treated with 200 µM concentration of both H_2_O_2_ and t-BHP as described earlier. The CypA levels in the conditioned media were quantified by using ELISA kit (Thermofisher) following the manufacturer’s instructions.

### Immunoblotting of CypA

In brief, 25 µg of protein extracts from H_2_O_2_/t-BHP treated and control cells were subjected to 12% SDS–PAGE. Following blotting of proteins, the nitrocellulose membrane was incubated in blocking buffer (tris-buffered saline containing 0.5% Tween-20 and 5% non-fat dried milk) for 1 hour at room temperature, followed by incubation with mouse monoclonal antibodies to cyclophilin A (1:1000), ß-actin (1:1000) and Nrf-2 (1:1000) for 2 hours at room temperature. The membrane was incubated with secondary goat anti–mouse IgG HRP antibody (Sigma) at a concentration of 0.2 µg/mL for 1 h at room temperature and the proteins were visualized using 3,3’-diaminobenzidine (DAB) reagent (Sigma) and H_2_O_2_ (Sigma). The densitometric analysis of the protein bands was performed using Image lab software.

### Estimation of somatic cell count and catalase activity in milk samples

Milk samples from various breeds of cattle such as Gir, Kankrej, Holstein Freisian (HF), white Holstein Freisian (WHF) and native breed of Gujarat were freshly collected to estimate the number of somatic cells. The milk samples (5 ml) were centrifuged at 12000 rpm for 15 minutes at 4°C, and the pellet was resuspended in 1 mL PBS. Added 0.4% of the trypan blue to 2µl of cell suspension and counted the total number of somatic cells using a cell counting machine. The enzymatic activity of catalase in milk samples was determined using catalase assay as described earlier in this study.

### Quantification of CypA levels in the milk samples

The levels of CypA protein in milk samples were quantified by using ELISA (Thermofisher) following the kit guidelines.

### Statistical analysis

All the experiments in this study were performed in triplicates. The statistical analyses were performed using the software, Graphpad, version 8.4.3. Data were compared by ordinary one-way ANOVA or two-way ANOVA, depending on the experimental design. Statistical significance was defined for p-values less than or equal to 0.05 (*), 0.01 (**), 0.001 (***), and 0.0001(****).

## RESULTS

### Culturing of mammary epithelial cells

Three different cell lines; GMECs, GLECs, and MMECs were used in the study. GMECs were successfully isolated from goat mammary glandular tissue, and a primary culture was established. The culture initially displayed a heterogeneous population of epithelial and fibroblast cells. Following differential trypsinisation, the epithelial cells cultured separately in growth medium. By the third day, cells formed a monolayer and exhibited typical cobblestone-like appearance of epithelial cells. To establish GLEC culture, primary epithelial cells isolated from fresh goat milk were seeded onto culture plates. A heterogeneous population of cells was found in the plates after 24 hours. After the end of third day, cells displayed a monolayer, cobblestone, epithelial-like morphology, characteristic to mammary epithelial cells. There were no notable differences in mammary epithelial cells isolated from mammary tissue and milk samples of goat. MMECs also exhibited a growth pattern with a typical epithelial phenotype by the end of third day of culture, but the cells were tightly packed when compared to those of GMECs and GLECs (Figure 1A).

**Figure 1.**
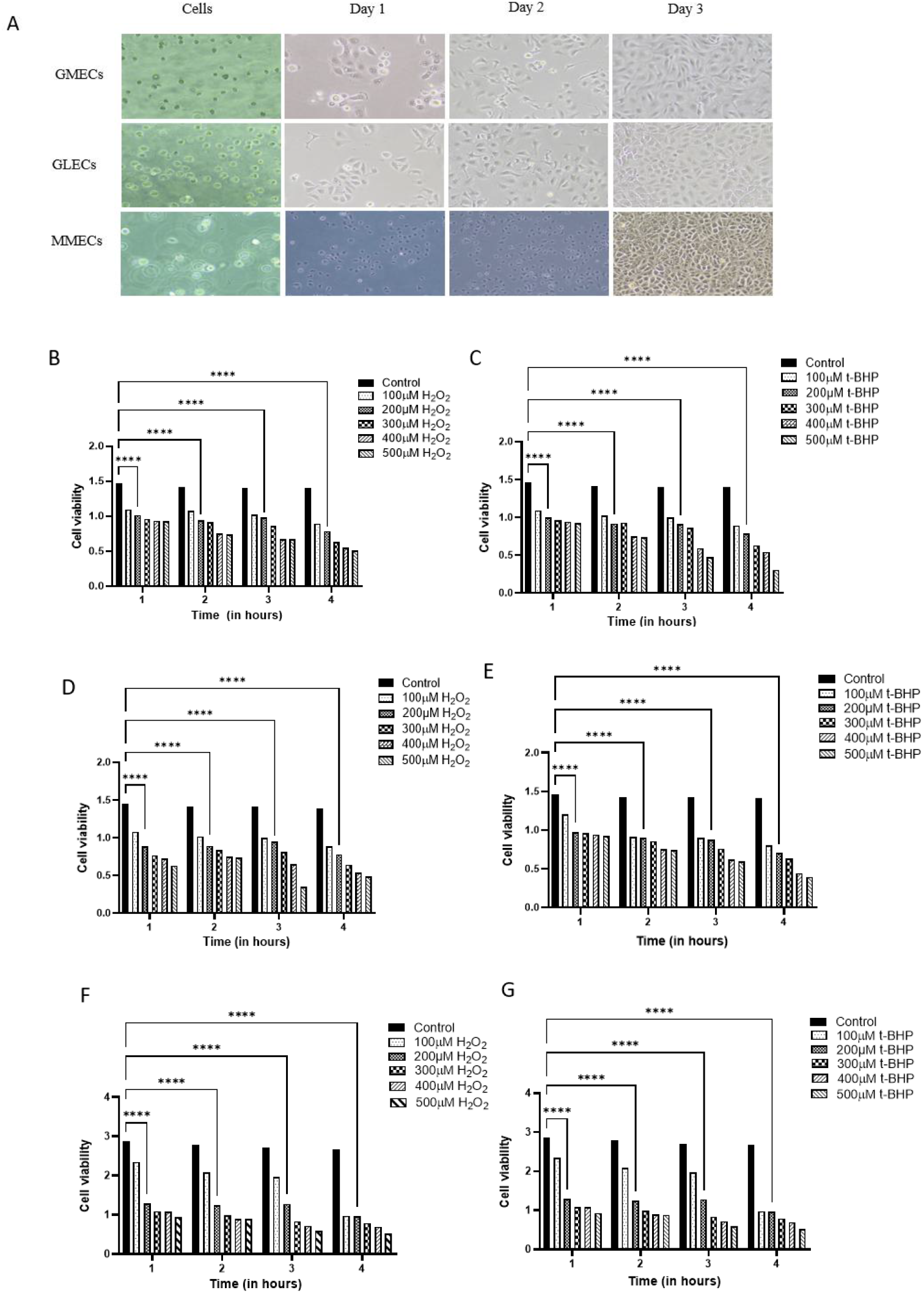
Mammary epithelial cell cultures of goat and mouse origin from day 1 to day 3 (A) Goat mammary epithelial cells (GMECs), goat luminal epithelial cells (GLECs), and mouse mammary epithelial cells (MMECs) in culture. Images captured using an inverted phase contrast microscope; 10x magnification. Relative viability of cells treated with 100 μM, 200 μM, 300 μM, 400 μM, and 500 μM concentrations of hydrogen peroxide (H_2_O_2_) and tert-butyl hydroperoxide (t-BHP), for a duration of 1 to 4 hours, was determined using MTT assay. (B & C) GMECs treated with H_2_O_2_ and t-BHP (D & E) GLECs treated with H_2_O_2_ and t-BHP, (F & G) MMECs treated with H_2_O_2_ and t-BHP. Statistical difference of p<0.05 was considered significant.

### Measurement of oxidative stress in the mammary epithelial cells

MTT assay was used to determine the viability of cells following treatment using H_2_O_2_ and t-BHP in varying concentrations and exposure time. Absorbance of the coloured product formed on treatment with 200 µM H_2_O_2_ and t-BHP in GMECs, GLECs, and MMECs was significantly different from that observed in control cells (p<0.0001) (Figure 1; B-G). Cellular viability of ∼70% was maintained in cells treated with 200 µM H_2_O_2_ and t-BHP for 1 hour. The cells exhibited a loss in viability when treated with >200 µM concentration and prolonged exposure of >1 hour. Hence, H_2_O_2_/t-BHP concentration of 200 µM with an exposure duration of 1 hour was considered as optimum for inducing oxidative stress in the cell types studied. Therefore, all further experiments were carried out with cells treated under these conditions.

### ROS levels were increased in mammary epithelial cells

One of the major indicators used for the measurement of ROS level is the fluorescent dye, DCF-DA, which is oxidized by ROS present in the cells *in vitro*, resulting in the formation of a fluorescent compound. The green fluorescence in H_2_O_2_/t-BHP-treated cells was significantly increased compared to non-induced cells, which indicated the presence of ROS, suggestive of a higher amount of oxidative stress (Figure 2; A). GMECs exhibited the highest intensity (p<0.001) when compared to that of the control group, followed by GLECs (p<0.01) and MMECs (p<0.05) on treatment with both the oxidative stress inducing agents (Figure 2; B, C, & D).

**Figure 2.**
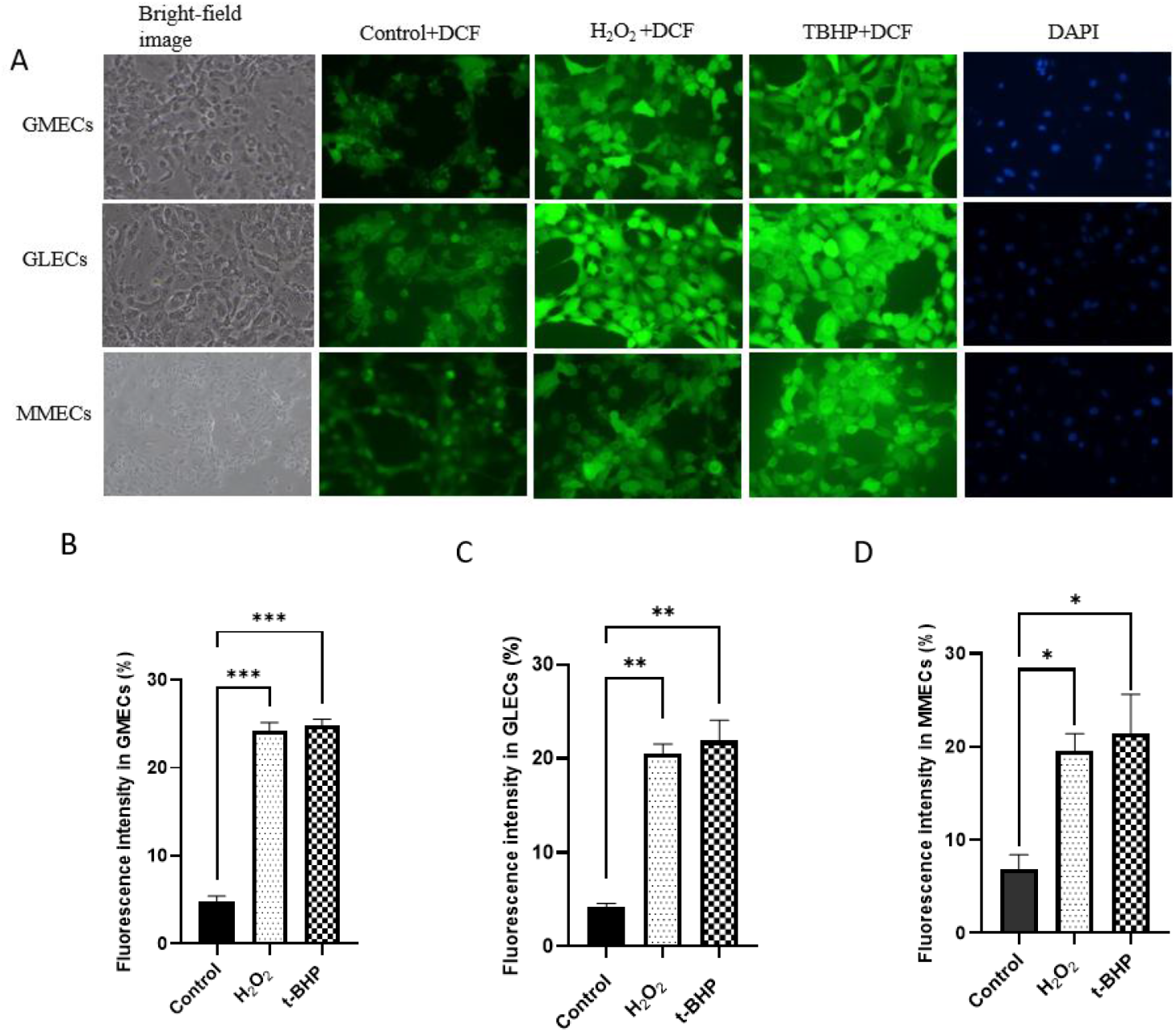
Measurement of intracellular ROS levels using DCF-DA (2′,7′-dichlorodihydrofluorescein diacetate; DCF-DA) and DAPI (4’,6-Diamidino-2-phenylindole) staining in mammary epithelial cells Representative images of goat mammary epithelial cells (GMECs), goat luminal epithelial cells (GLECs), and mouse mammary epithelial cells (MMECs) with and without treatment (control) with hydrogen peroxide (H_2_O_2_) or tert-butyl hydroperoxide (t-BHP). Percentage fluorescent intensity of experimental groups (B) GMECs, (C) GLECs and (D) MMECs. Data represented as mean ± SD. Statistical significance was defined for p-values less than or equal to 0.05 (*), 0.01 (**), and 0.001 (***).

Lipid peroxidation index in the cell cultures was determined by using TBARS assay. In this experiment, the levels of malondialdehyde produced as the end product of lipid breakdown were compared in the cell types. A marked increase in lipid peroxidation after incubation with pro-oxidants, implicated that the mammary epithelial cells are extremely sensitive to oxidative damage. The rate of malondialdehyde production was significantly higher in H_2_O_2_/t-BHP treated cells with a p-value of 0.0001 for GMECs, and p<0.001 for both GLECs and MMECs in comparison to the untreated cells (Figure 3; A, B & C).

**Figure 3.**
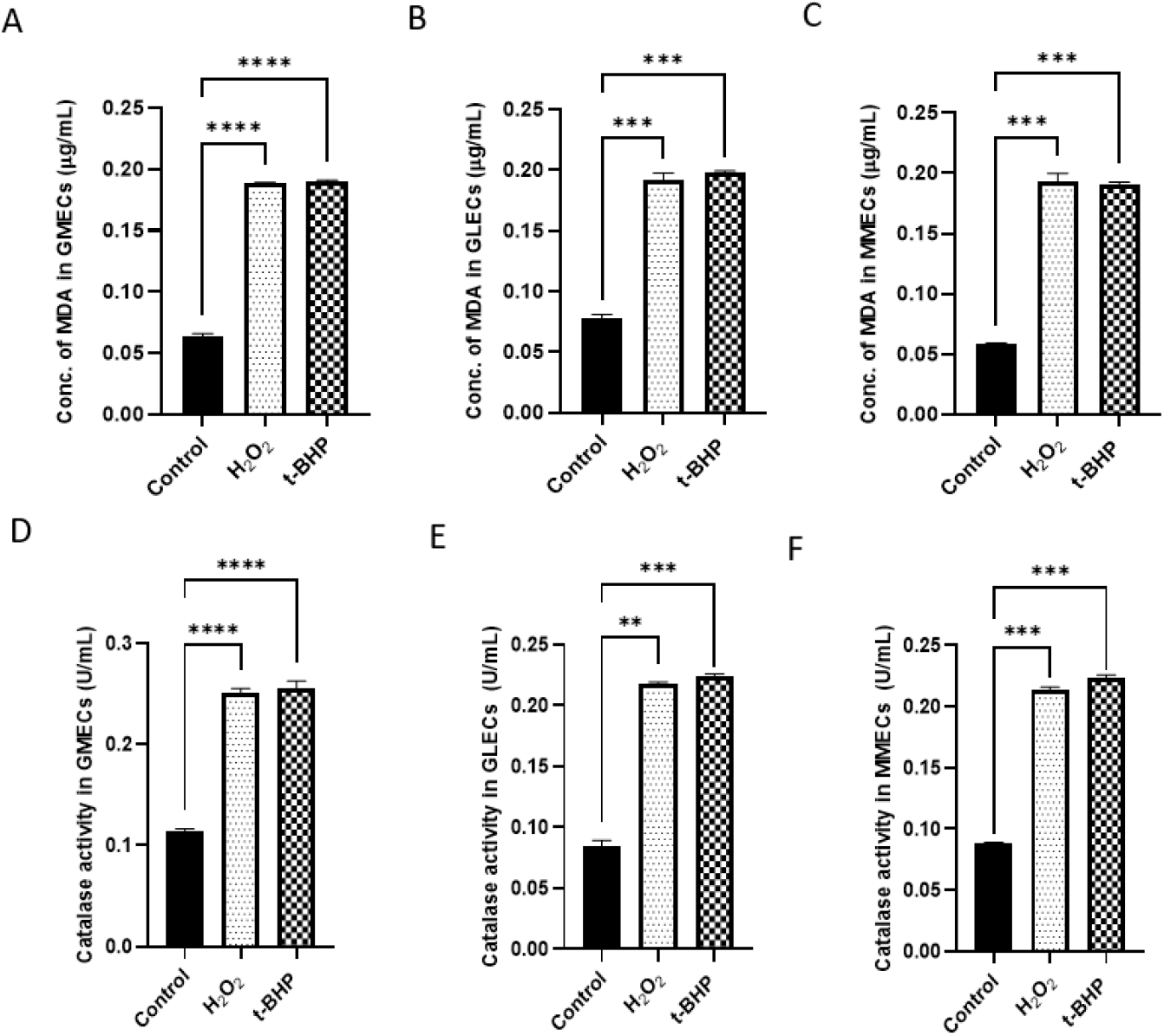
Graphical representation of lipid peroxidation index and catalase enzyme activity in goat mammary epithelial cells (GMECs), goat luminal epithelial cells (GLECs), and mouse mammary epithelial cells (MMECs) treated with hydrogen peroxide (H_2_O_2_) or tert-butyl hydroperoxide (t-BHP). Levels of malondialdehyde in (A) GMECs, (B) GLECs and (C) MMECs. Enzymatic activity of catalase in (D) GMECs, (E) GLECs and (F) MMECs. Data represented as mean ± SD. Statistical significance was defined for p-values less than or equal to 0.01 (**), 0.001 (***), and 0.0001(****).

Catalase is an antioxidant enzyme that catalyzes the decomposition of H_2_O_2_ into water and oxygen, hence decreasing the level of oxidative stress in the cell. Our results demonstrated that there was a significant increase in total catalase activity in the cells treated with either H_2_O_2_ or t-BHP as compared to the untreated cells. Among the cell types studied, the catalase activity of the GMECs was significantly higher than that of the untreated group (p<0.0001) and there was no notable difference in enzymatic activity when treated with both H_2_O_2_ and t-BHP (Figure 3; D). The GLECs exhibited a significantly higher catalase activity when treated with t-BHP (p<0.001) than when treated with H_2_O_2_ (p<0.01) in comparison to control group (Figure 3; E). MMECs, exhibited a comparable catalase activity on treatment with both the oxidising agents when compared to the control group (p<0.001) (Figure 3; F).

### Expression of CypA in mammary epithelial cells

The Cyp A secreted from mammary epithelial cells with and without H_2_O_2_ or t-BHP treatment was quantified using ELISA. The levels of CypA in all the cell types studied were approximately 0.3ng/mL on treatment with both the oxidising agents. A two-fold increase of CypA was observed in the conditioned media of GMECs treated with H_2_O_2_ or t-BHP than that of the control group with a p-value of 0.001 (Figure 4; A). A significant increase in the CypA levels was also noticeable in the other two cell types in comparison to the control group (p<0.0001) (Figure 4; B & C).

**Figure 4.**
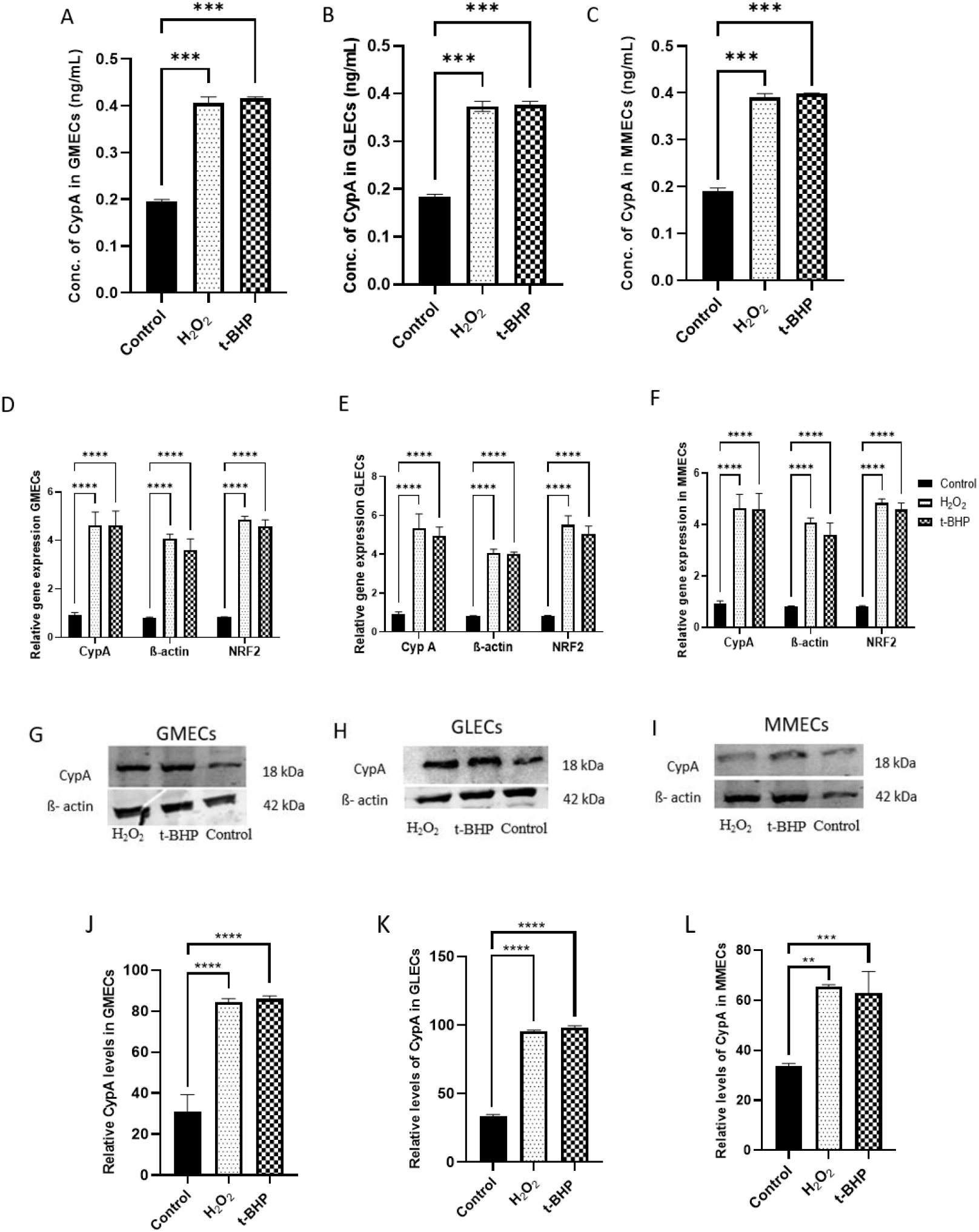
Cyclophilin A levels in goat mammary epithelial cells (GMECs), goat luminal epithelial cells (GLECs), and mouse mammary epithelial cells (MMECs). Quantitation of cyclophilin A in the conditioned media of (A) GMECs, (B) GLECs, and (C) MMECs by ELSIA. Gene expression levels of cyclophilin A in (D) GMECs, (E) GLECs, and (F) MMECs. Representative immunoblots and relative abundance of cyclophilin A protein in (G & J) GMECs, (H & K) GLECs, and (I & L) MMECs. Data represented as mean ± SD. Statistical significance was defined for p-values less than or equal to 0.05 (*), 0.01 (**), 0.001 (***), and 0.0001(****).

### CypA gene expression in mammary epithelial cells

The expression of *cypA* gene was quantified using qRT-PCR in H_2_O_2_ and t-BHP-treated cells and compared with the control cells. The transcription factor NRF2 was used as a positive control. NRF-2 is known to regulate cellular defence at the transcriptional level in the case of oxidative stress. ß-actin was used as an endogenous control for normalizing the expression levels of *cypA* gene. The expression levels of *cypA* were significantly higher in all the cell types treated with H_2_O_2_/t-BHP when compared to that of the untreated cells (p<0.0001). The data also suggested that the expression was highest in GMECs followed by GLECs and MMECs (Figure4; D, E, & F). It indicates differential responsiveness and sensitivity towards oxidative agents among these cell lines.

### CypA protein expression in mammary epithelial cells

The expression of CypA protein in GMECs, GLECs, and MMECs under oxidative stress and control conditions was investigated using western blotting technique. Immunodetection revealed the presence of a band at 18 kDa region that corresponds to the molecular weight of CypA protein (Figure 4; G, H, I). The relative abundance of CypA protein was significantly increased in GMEC and GLEC cells (p<0.0001) on treatment with H_2_O_2_/t-BHP (Figure 4; J, & K). In MMECs, there was a modest increase in expression (p<0.01) when compared to untreated cells (Figure 4; L).

### Somatic cell count and CypA levels in cattle milk

The somatic cell count in the milk samples collected from WHF, Kankrej, HF, Gir and native breeds of dairy cows are represented in Table 2. The cell count was observed to be highest in the milk sample obtained from Kankrej breed (1.91×10^7^ cells/mL) whereas the HF breed showed the least number of cell count (1.33×10^5^ cells/mL). The catalase activity in milk samples from Kankrej breed, where somatic cell numbers were relatively high, displayed the highest enzymatic activity when compared to the milk from other breeds of cow (Figure 5; A). The CypA levels in the milk samples were observed to be in a range of 1.5-2.05 ng/mL (Figure5; B).

**Table 2:**
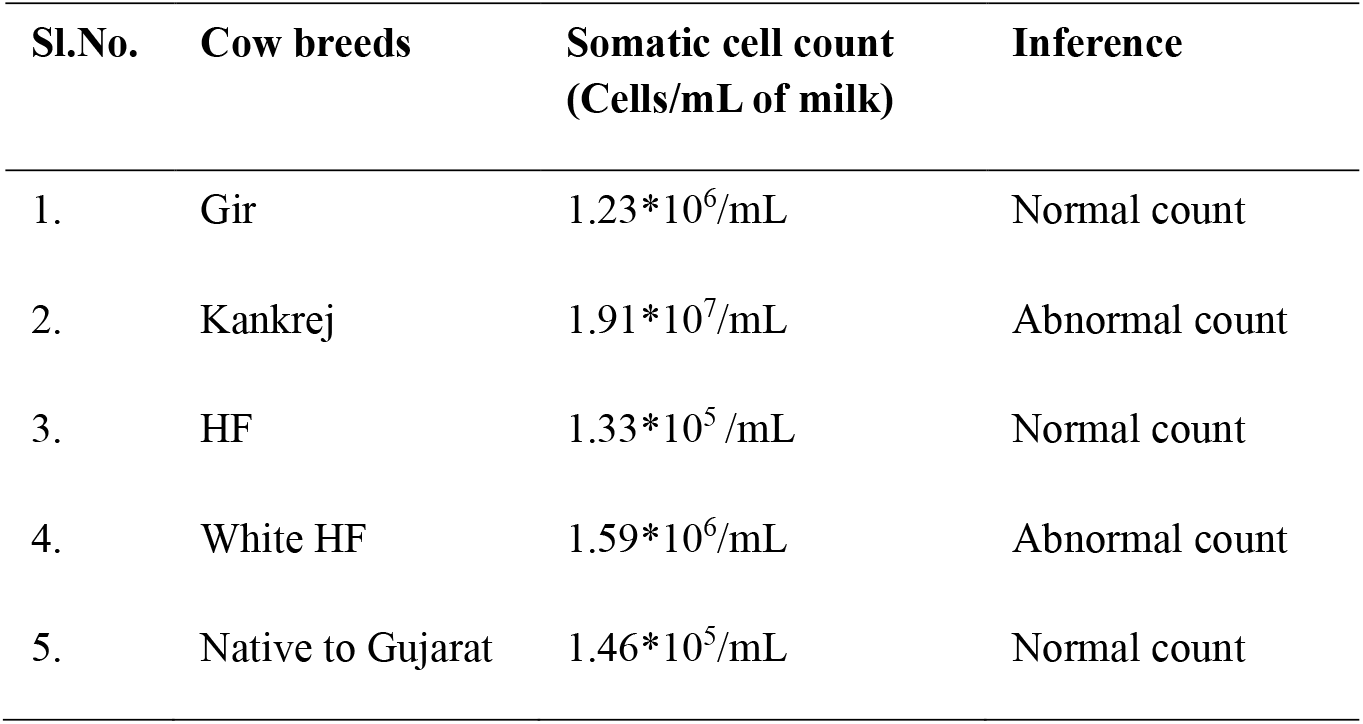
Somatic cell count in the milk samples of various breeds of dairy cows.

**Figure 5.**
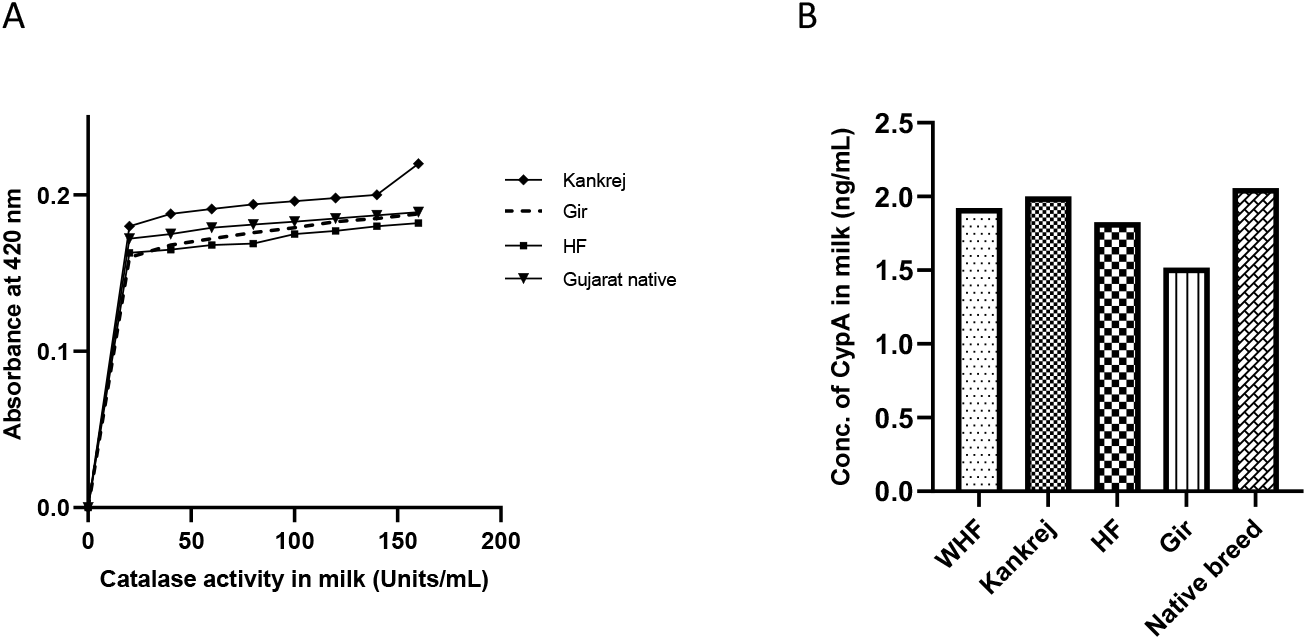
Fig 5: Analysis of milk samples from various breeds of cow (white Holstein Freisian (WHF), Kankrej, Holstein Freisian (HF), Gir, and native breed of Gujarat (A) Representative graph showing the enzymatic activity of catalase in milk samples and Quantification of CypA (Cyclophilin A) in milk samples from various breeds of dairy cows.

## DISCUSSION

Here, we report CypA as a redox-sensitive protein, secreted by mammary epithelial cells of goat and mice. We also demonstrate that CypA is detectable in the milk of dairy cows, and it could serve as an effective oxidative stress-induced secretory marker in milk.

Despite being one of the highest milk-producing countries, India has shown a decline in milk production due to various kinds of stress including excessive heat, humidity, pollution, and climatic changes that adversely affect livestock milk production [12]. Dairy cows undergo numerous transitional phases, like periparturient and neonatal stages, which make them prone to various metabolic and infectious diseases [9]. All these factors that compromise the efficiency of cattle may be due to oxidative stress resulting in free radical generation that promotes inflammation of mammary gland, and the milk production is largely affected [13]. The dairy sector in India plays an important role in nutritional security, sustenance and development of economy [14]. Hence, the oxidative stress management in dairy cows is a crucial issue that needs to be addressed. The quantification of antioxidant enzymes such as superoxide dismutase, glutathione peroxidase, catalase activity etc., in the blood have been reported to assess the oxidative status in animals however, there is no established biological marker for assessing oxidative stress [11].

CypA is an intracellular protein that is secreted in response to inflammatory stimuli [15]. Studies have shown that CypA is involved in various cellular functions such as protein folding, trafficking and cell signalling [16]. In humans, it has been implicated in several pathological conditions such as cardiovascular, neurodegenerative and immune disorders, viral infections, cancer, etc. [17]. The secreted form of CypA functions as a chemokine that mediates intercellular communication and has been shown to stimulate proinflammatory signals in endothelial cells, vascular smooth muscle cells, and leukocytes [18]. However, the role of CypA in regulating oxidative status in animals or in mammary epithelial cells has not been explored.

In the present study, we first established an *in vitro* oxidative stress model in mammary epithelial cells (GMECs, GLECs, and MMECs) by treating the cells with oxidative stress-inducing agents like H_2_O_2_ and t-BHP. A concentration of 200 µM of both H_2_O_2_ and t-BHP and an exposure duration of 1 hour was observed to be optimum in inducing oxidative stress, with which the cells were viable without causing excessive damage. H_2_O_2_ is a naturally produced oxidizing agent that acts as a direct ROS and is reported to induce oxidative stress in mammary epithelial cell lines [19]. t-BHP is a synthetic drug that generates free radicals through metabolic pathways involving cytochrome-P450 and glutathione peroxidase and causes oxidative damage to the cells. Both H_2_O_2_ and t-BHP are known to deplete cellular antioxidant defense mechanisms, initiate lipid peroxidation and reduce mitochondrial membrane potential in *in vitro* systems [20]. Huang et al., 2024 have studied the protective effects of hesperidin, an anti-inflammatory flavonoid, in alleviating the harmful effects of oxidative stress, such as ROS production using H_2_O_2_ treated bovine mammary epithelial cells (BMECs) [21].

Mammary epithelial cells are located on the apical surface of the alveoli and the ducts of mammary glands in cows. Inspite of their role in milk production, mammary epithelial cells also function as the first line of immune defence in mammary gland of dairy cows. Hence, mammary epithelial cell lines have been widely used to study oxidative stress and inflammatory responses in dairy cows [22]. Oxidative stress triggers the activation of redox-sensitive signaling pathways, which initiate the transcription of proinflammatory genes, such as IL-1β, IL-6, and TNF-α [23]. It also induces caspase signaling cascade in mammary epithelial cells, leading to cell death and deterioration of mammary gland [24]. Zhuang et al., 2020, has used bovine mammary epithelial cells as a model of oxidative stress to study immune responses triggered by *E*.*coli* infections [25]. Transcriptome profiling of bovine mammary epithelial cells challenged by CypA, *Escherichia coli* derived lipopolysaccharide and heat-killed strains of *Staphylococcus aureus* for inducing oxidative stress have been used to study pathways associated with inflammatory responses. This study also reveals the potential of CypA to trigger the release of various cytokines in these cell lines [26]. BMECs have been widely used to study bovine mastitis, which is the most contagious and infectious disease that affects dairy cows. *E. coli* and *S. aureus* are the most prevalent pathogens responsible for causing mammary gland infection. Mastitis can affect milk composition, its quality as well as the quantity, which accounts for huge economic loss [27].

We measured oxidative stress in the cell lines by using standard protocols of DCF-DA staining, TBARS assay, and catalase assay. The results of DCF-DA assay showed a significantly increased level of ROS production in all three cell lines by treatment with 200 µM concentration of both t-BHP and H_2_O_2_ as indicated by the fluorescence intensity. The densitometric analysis showed that the intensity was more pronounced in the GMEC cell lines followed by the GLECs and MMECs. We found that incubation of GMECs cells with 200 μm H_2_O_2_/t-BHP, significantly damaged the cells with altered cell morphologic appearance, decreased cell viability, and increased ROS levels. DCF-DA assay provides a direct measurement of the cellular ROS levels. DCF-DA is fluorescent probe that diffuses into the cells gets hydrolyzed by intracellular esterases which finally reacts with H_2_O_2_, to generate a fluorescent product, 2′,7′-dichlorofluorescein. Therefore, the fluorescent intensity generated in the assay corresponds to the amount of peroxide produced by the cells [28]. Intracellular ROS production in BMECs challenged with a mycotoxin was determined in a study using DCF staining assay in which an increased level of fluorescent intensity was observed in the cells exposed to toxin versus the control cells [29]. Another study by Castellani et al, 2024 has also shown that the fluorescent intensity produced by the DCF assay is directly proportional to the accumulation of ROS in BMECs subjected to acute temperatures [30].

Lipid peroxidation levels were tested in the mammary epithelial cells, revealing increased production of malondialdehyde in GMECs isolated from the mammary gland of goat. ROS like free radicals can interact with lipids in the cell membrane, causing oxidative damage and results in the formation of lipid peroxidation products like malondialdehyde. In TBARS assay, the malondialdehyde reacts with thiobarbituric acid at high temperatures in acidic conditions to form a pink-colored substance which when quantified gives an index of lipid peroxidation. Jiang et al, 2022 has investigated the protective effects of flavonoids on inflammation responses and oxidative stress, Lipopolysaccharide (LPS) induced BMECs [31]. Elevated levels of malondialdehyde were observed in the cells in response to oxidative stress induction, which is consistent with our findings. We next tested for catalase activity in the cellular models treated with t-BHP and H_2_O_2_. Catalase is a ubiquitous antioxidant enzyme that is present in almost all body tissues and plays an important role in alleviating oxidative stress by catalyzing the decomposition of H_2_O_2_ into water and oxygen [32]. We found an increase in catalase activity in all three cell lines post-treatment with H_2_O_2_/t-BHP which demonstrated the ability of mammary epithelial cells to respond to redox stress.

The CypA levels in the conditioned media of the cultured mammary epithelial cells with and without H_2_O_2_/t-BHP treatment was also quantified by ELISA. The CypA levels were significantly higher in the conditioned medium obtained from the stress-induced cells (0.3 ng/mL) versus untreated cells (0.12 ng/mL). This is the first report wherein CypA levels have been quantified from mammary epithelial cells of goat and mouse origin.

The transcript level expression of CypA were also assessed in the study using RT-PCR analysis. It was observed that the gene expression of CypA was 5-6 fold higher in the H_2_O_2_/t-BHP treated cells as compared to that of the untreated cells. There was no significant difference in the CypA expression levels with both H_2_O_2_ and t-BHP treatment. Our study thus validated the presence of an intracellular form of CypA in mammary epithelial cells. Immunoblotting of CypA in the control and treated cell lysates further confirmed the presence of CypA protein at 18 kDa region which is similar to the previous reports [8]. The relative levels of CypA was significantly higher in the lysates from H_2_O_2_/t-BHP treated samples in comparison to the untreated samples. The immunoblotting data demonstrated that both types of goat mammary cell activate CypA as part of their response to oxidative stress in comparison to mouse mammary epithelial cells. This refers to the species-specific responsiveness to the expression of CypA to oxidative stress. Our study utilizing cell lines from two different species indicates that the role of extracellular CypA is highly conserved across species.

Further, we observed that catalase activity is increased in milk samples collected from different breeds of dairy cows from villages in Gujarat, indicating that the animals were exposed to oxidative stress. An elevated somatic cell count of diluted milk also implicated the presence of bacterial infection in the milk samples. The somatic cell count in the milk samples is an indicator of udder health and milk quality in dairy cows. There is no established standard cut-off for somatic cell count in India [33]. A somatic cell count of <2 lakh/ mL is generally considered normal and an elevation of cell count to >2 lakh cells/mL suggests an inflammatory response in the mammary gland of animal and an increased susceptibility to infection [34]. A study by Sarvesha et al., 2017 has reported that SCC of >5 lakhs/mL of milk indicates the presence of bacterial infection in the milk samples [35]. Among the samples analysed in our study, milk obtained from WHF and Kankrej breeds had SCC of >5 lakhs/mL. The CypA levels in the milk samples were observed to be in a range of 1.5 -2.05 ng/mL. It is also noted that these levels were twice higher than those quantified in the conditioned medium. A recent study by Takanashi et al, 2025, has reported significantly higher concentrations of CypA in dairy cows subjected to intramammary infusion of mastitis-causing *S. aureus* bacteria when compared to that of healthy dairy cows [8]. They have also reported a significant positive correlation with the SCC and CypA levels in milk. A previous study from the same research group has reported the extracellular secretion of CypA in the mammary glands of cattle with mastitis and have shown the chemotactic potential of CypA to recruit granulocytes to the site of inflammation. They have also demonstrated the anti-bovine CypA antibody could also inhibit inflammatory responses mediated by extracellular CypA [36].

There is possibly an important role for CypA in the cellular response to oxidative stress (Figure 6). Further studies need to be conducted to understand the mechanisms by which CypA modulates mammary cell function during oxidative stress and its subsequent effects on milk production in cattle.

**Figure 6.**
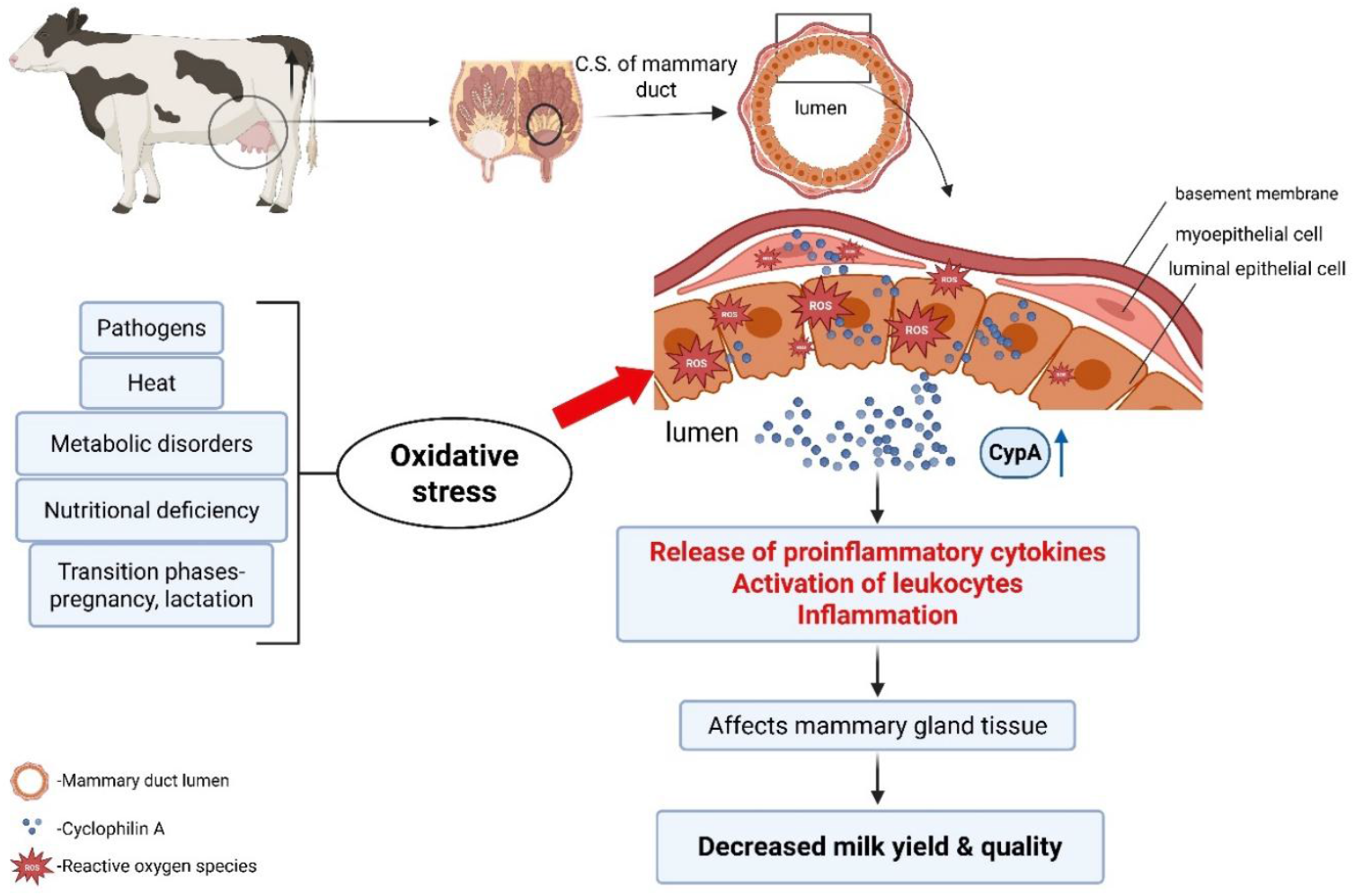
Cyclophilin A (Cyp A) secretion in response to oxidative stress. CypA is secreted by epithelial cells that lines the alveoli and lumen of mammary duct in the udder of dairy cows, in response to oxidative stress induced by various factors such as pathogenic infections, heat stress, metabolic disorders, nutritional deficiencies, pregnancy and lactation. Cyp A triggers inflammatory response which negatively affects the mammary gland tissue resulting in decreased milk yield and quality. (Created with BioRender.com).

## CONFLICT OF INTEREST

None

## Notes

### Competing Interest Statement

The authors have declared no competing interest.

